# Modulation of β-catenin levels is critical for cranial neural crest patterning and dispersal into first pharyngeal arch

**DOI:** 10.1101/726299

**Authors:** Alok Javali, Vairavan Laxmanan, Dasaradhi Palakodeti, Ramkumar Sambasivan

## Abstract

Vertebrate cranial neural crest cells (CNCC) are multipotent. Proximal to the source CNCC form the cranial ganglia. Distally, in the pharyngeal arches, they give rise to the craniofacial skeleton and connective tissues. Fate choices are made as CNCC pattern into distinct destination compartments. In spite of this importance, the mechanism patterning CNCC is poorly defined. Here, we report that a novel β-catenin-controlled switch in the cell arrangement is critical in patterning CNCC. In mouse embryos, at the first pharyngeal arch axial level, membrane β-catenin levels correlate with the extent of cell-cell adhesion and thus, with a collective or a dispersed state of CNCC. Using *in vitro* human neural crest model and chemical modulators of β-catenin levels, we show a requirement for down-modulating β-catenin for the collective-to-dispersed switch. Similarly, in β-catenin gain of function mutant mouse embryos, CNCC fail to disperse, which may underlie their failure to populate first pharyngeal arch. Thus, we show that β-catenin-mediated regulation of CNCC tissue architecture, a previously underappreciated mechanism, underlies the patterning of CNCC into fate-specific compartments.

**Summary statement:** The report shows a crucial step in cranial neural crest patterning. Neural crest cells invading the pharyngeal arches transition from a collective to a dispersed state. This transition in cell arrangement is dependent on membrane β-catenin levels.

## Introduction

The ectoderm-derived cranial neural crest cells (CNCC) have extraordinary differentiation potential and are central to the formation of the vertebrate head (Gans and Northcutt, 1983; Couly, Coltey and Le Douarin, 1993). CNCC contributes to the formation of neurons and glia of the cranial ganglia, pigment cells, dermis as well as craniofacial skeleton (Le Douarin *et al.*, 2004; Cordero *et al.*, 2010; Bhatt *et al.*, 2013). Broadly, the anatomical location of post migratory CNCC correlates with their fate. The cells contributing to craniofacial skeleton arise from a distinct spatiotemporal domain in the neural plate border as opposed to those contributing to the neural or pigment lineage, suggesting divergence of fate at origin (Raymond Teck Ho Lee *et al.*, 2013; Weston and Thiery, 2015). However, recent evidence tracing developmental trajectory using single cell transcriptomics demonstrate that fate decisions are made in a stepwise fashion as CNCC migrate and pattern (Soldatov *et al.*, 2019). This highlights the role of signalling cues *en route* and at the destination in the fate restriction of CNCC lineage. Thus, the patterning of CNCC is tightly linked with its normal ontogeny.

CNCC emerges from lateral edges of the anterior neurectoderm, where they undergo epithelial to mesenchymal transition and begin to migrate extensively in stereotypic manner (Kulesa and Gammill, 2010; Theveneau and Mayor, 2012; Bhatt *et al.*, 2013; R. T. H. Lee *et al.*, 2013). Upon delamination, the directionality of CNCC migration is determined as a result of interactions with neighbouring tissues and the signal rich microenvironment they face. Delaminated cells in the hindbrain migrate collectively in three distinct streams to invade different pharyngeal arches. Secretion of neural crest repelling molecule such as Semaphorin-3A, prevents mixing between the distinct migratory streams of cells by creating neural crest free zone between the streams, thus, orchestrating patterning of CNCC derived tissue in rostro-caudal axis (Kulesa *et al.*, 2010). However, within each of the migrating streams, mechanism of patterning CNCC into distinct fate specific domains is unclear.

Two independent computational models, supported by experimental evidence, address the directionality of migratory CNCC stream, which relies on chemotaxis driven by molecules, VEGF and SDF1 (Theveneau *et al.*, 2010; McLennan *et al.*, 2015; Szabó and Mayor, 2016). Notably, both models highlight the importance of collective behaviour mediated by cell-cell adhesion in sensing the chemo-attractants. Two distinct phases of migration is seen in CNCC emerging from xenopus embryo explants. In the initial phase, reminiscent of chemotaxis-mediated early migratory CNCC *in vivo*, explant-derived cells maintain stable cell-cell contacts and migrate as a sheet. In the second phase, cells are dispersed and display more mesenchymal morphology (Cousin, 2017). CNCC arising from mouse embryo explants also display these two distinct tissue architecture (Gonzalez Malagon *et al.*, 2018). Remarkably, transitions in tissue arrangement drive key morphogenetic events during vertebrate embryogenesis (Thiery, 2003; Mongera *et al.*, 2018). However, such collective / dispersed state transition and its relevance in the patterning of CNCC *in vivo* has not been addressed.

In this study, using mouse embryos, we investigated the tissue-level architecture of migratory and post-migratory CNCC at the level of first pharyngeal arch. We find that CNCC retain stable cell-cell contact mediated by N-cadherin following epithelial to mesenchymal transition and stay as a cluster at the proximal location, where it forms the trigeminal ganglia. However, distally migrated cells, which invade pharyngeal arches and generate mandible, dermis and connective tissues, manifest a dispersed organisation. Reduced membrane β-catenin levels strongly correlate with collective to dispersed transition of CNCC and exogenous modulation of β-catenin levels effect similar transition in human neural crest model. Moreover, by revisiting previously reported neural crest specific β-catenin gain of function mutant, we reveal that the failure of CNCC to invade first arch is associated with the failure of dispersal of the distal pool. Together, our findings show a novel role for β-catenin-mediated regulation of cell arrangement in patterning CNCC into fate specific compartments.

## Results and Discussions

### Collective and dispersed states of cranial neural crest correlate with membrane β-catenin levels

Cranial neural crest from both Xenopus and mouse embryo explants display collective as well as dispersed cell arrangement. To test whether such distinct tissue arrangement is present *in vivo* and across species, we analysed CNCC in mouse embryos and in neural crest like cells (NCLC) derived from human embryonic stem cells (Bajpai *et al.*, 2009). Derivation of NCLC involves induction of neurulation from embryonic stem cells to obtain self-organized structures called neurospheres. Upon induction of attachment on Fibronectin-coated plates, migratory cells emerge from the neurospheres, which express a wide panel of neural crest markers (Fig. 1A, B, S1). The migratory cells are NCLC, which could be enzymatically dissociated and passaged. NCLC express several cranial neural crest specific genes as revealed by comparative transcriptome analysis between NCLC and neurospheres prior to attachment (Fig. S2). NCLC also exhibit extraordinary differentiation potential as they can be differentiated into neural as well as skeletal lineages (Bajpai *et al.*, 2010).

**Figure 1:**
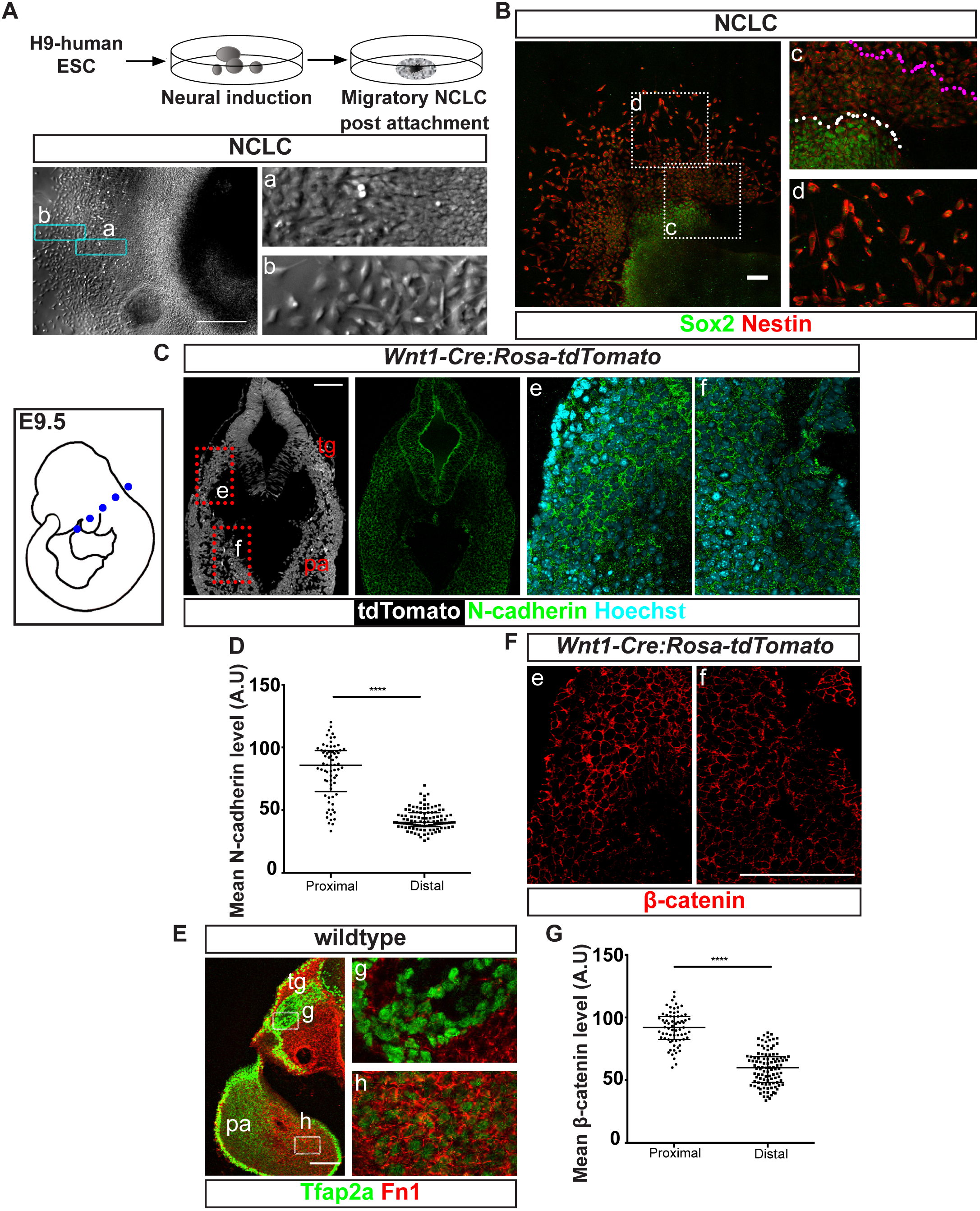
β-catenin levels in cell membrane correlate with differential cell arrangements in cranial neural crest cells. (A&B) Migratory NCLC emerging from neurospheres derived from human embryonic stem cells as seen in bright field (A) or immunostained to detect Sox2 (neural) and Nestin (neural crest) markers (B). Zoomed in image on the right: NCLC proximal (a and c) to the attached spheres appear as a collective and distal NCLC (b and d) display dispersed morphology. White dotted line demarcates neurospheres and NCLC, magenta line demarcates proximal and distal NCLC. (C) Transverse section of E9.5 *Wnt1-Cre* based neural crest reporter embryo at the level of first pharyngeal arch showing reporter expression (Left) and N-cadherin expression (Right). Extreme right panels are zoomed in images of N-cadherin staining. (D) Quantitation of N-cadherin levels in the membrane of tdTomato positive cells in proximal and distal regions. A.U. – arbitrary units. (E) Transverse section immunostained for Tfap2a and Fibronectin. Right panel shows zoomed in images of indicated regions g and h. Note the difference in Fn1 distribution between proximal (g) and distal (h) CNCC. (F) Images from the same optical section in C and corresponding to the same zoomed in regions e and f, showing β-catenin staining. Note the correlation in the levels of N-cadherin and β-catenin. (G) Quantitation of β-catenin expression in the membrane of tdTomato positive cells in proximal and distal regions. tg-anlage of trigeminal ganglia, pa-pharyngeal arch. Statistical analysis: two tailed unpaired t test. **** is p<0.00001. Scale bar: 100μm

Here, we show that migratory NCLC emerging from neurospheres display morphologically distinct phases, similar to Xenopus and mouse explant cultures. Proximal to neurospheres NCLCs are tightly packed and distal cells are dispersed (Fig. 1A, B). To test whether such distinct architecture is relevant *in vivo*, we analysed post-migratory CNCC at the level of first pharyngeal arch in E9.5 mouse embryos. Transverse sections of *Wnt1-Cre:ROSA*^*tdTomato*^ neural crest reporter embryos were assayed for the expression of Ca^2+^ dependent cell adhesion molecule N-cadherin. We observed that proximally located anlage of trigeminal ganglia display abundant N-cadherin as opposed to reduced levels in the distal skeletogenic CNCC compartment in the 1^st^ pharyngeal arch (Fig. 1C, D, Fig. S3A). A gradient distribution was observed among CNCC within the pharyngeal arch as well, however, the difference between ganglia anlage and arch was still striking (Fig. S3B). Lower levels of N-cadherin suggest weak cell-cell junctions and dispersion in the distal domain. To directly assess dispersion, we assessed the distribution of extracellular matrix. Co-immunostaining was performed using antibodies to detect TFAP2a and Fibronectin 1 (Fn1), a key extracellular matrix component. The analysis revealed uniform distribution of Fn1 around individual cells of the CNCC pool in pharyngeal arches suggesting the presence of interstitial matrix (Fig. 1E). On the other hand, Fn1 staining was sparse around the clusters of cells in the proximal CNCC domain (Fig. 1E). Differential N-cadherin abundance and a concordant variation in Fn1 distribution indicate that the discrete proximodistal CNCC compartments are characterized by distinct cellular arrangement.

The difference in post-migratory CNCC organisation could result from two possibilities; A) selective switching of distal pool from collective to dispersed state or B) emergence of two distinct pools at the time of CNCC delamination from the neural plate border. To distinguish between these possibilities, we analysed a time series in mouse and human models. At 4 somite stage mouse embryos, the delaminated CNCC are found in a cluster, whereas at 8 somite stage, the migratory CNCC display a proximal collective and distal dispersed organisation (Fig. S4A). Similarly, NCLC that emerge at 12 hrs are a collective, while later at 24 hrs NCLC disperse distally (Fig S4B). Thus, distal pool destined to arches appear to disperse from a collective. Notably, similar transition in axial progenitors, from collective to dispersed arrangement underlies differential tissue fluidity, which acts as a major driver of embryonic morphogenesis (Mongera *et al.*, 2018). In the context of CNCC, the significance of this transition and the underlying molecular mechanism are unclear.

First, to investigate the mechanism, we looked at the distribution of β-catenin. In addition to Wnt-responsive transcription regulation in the nucleus, β-catenin in the cell membrane regulates adhesion. Membrane β-catenin is an important component of cadherin mediated cell-cell adhesion complex and acts to stabilise cell-cell contacts. (Brembeck, Rosário and Birchmeier, 2006; Nelson, 2008). β-catenin binds to actin cytoskeleton as well as the cytoplasmic domains of type-I cadherins, potentially causing clustering of cadherins on the membrane (Maher *et al.*, 2009). In CNCC, β-catenin expression was predominantly found in cell membrane. Strikingly, the distribution pattern of membrane β-catenin matched that of N-cadherin, with high levels in proximally located CNCC and reduced levels in skeletogenic CNCC in pharyngeal arches (Fig. 1F, G, S5). Interestingly, a recent study showed dynamic modulation of nuclear β-catenin levels in migratory CNCC following their induction in *Xenopus* embryos (Maj *et al.*, 2016). This prompted us to investigate the role of β-catenin modulation in regulating tissue architecture in CNCC compartments.

### Stabilisation of β-catenin promotes compaction of neural crest-like cell clusters

To test whether change in β-catenin levels regulates collective / dispersed behaviour of cranial neural crest, we employed human neural crest model. P0-NCLC *in vitro* were treated with pharmacological modulators of β-catenin stability, Chiron 99201 (referred to as Chiron) and XAV-939 (referred to as XAV) to stabilise and destabilise β-catenin, respectively (Fig. 2A). Upon Chiron treatment, TFAP2a+ NCLC in the outer periphery, i.e., located distally from the parent neurosphere, formed compact clusters instead of dispersing (Fig. 2B). Cells on the edge of individual clusters lacked lamellipodia, the actin-based membrane extensions associated with active migration, typically observed in dispersed control NCLC. This is consistent with previous reports that showed decreased spread of migratory neural crest cells from mouse or *Xenopus* neural crest explants treated with GSK3β inhibitors, BIO or Chiron (Maj *et al.*, 2016). Conversely, treatment with XAV led to the formation of patches of dispersed cells with pronounced mesenchymal morphology and increased actin stress fibres compared to vehicle control (Fig. 2B). The morphology and the architecture were not uniform in the outer periphery; however, they were unique to XAV treatment. This observation in human NCLC extends our correlative observation in mouse embryos and points to a causal role for changing β-catenin levels in collective versus dispersed state.

**Figure 2:**
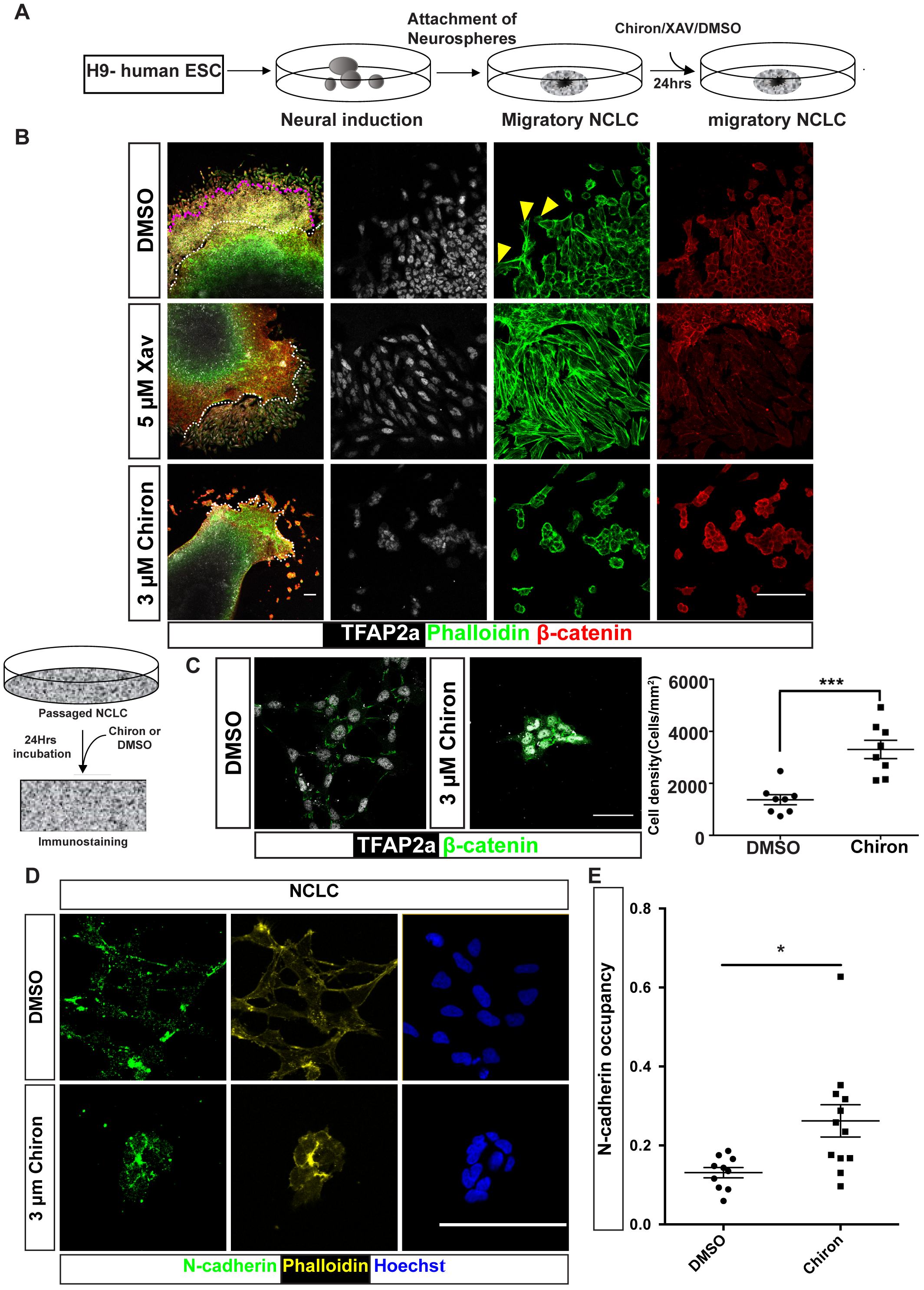
Stabilisation of β-catenin promotes cell cohesion and collective-like behaviour in human NCLC. (A) Schematic of the experiment for modulation of β-catenin levels in migratory NCLC. (B) Migratory NCLC treated with β-catenin modulators or vehicle control, immunostained to detect β-catenin and TFAP2a; actin was stained with phalloidin to assess cell morphology. Yellow arrow heads mark lamellipodia. White dotted line demarcates neurospheres and NCLC, magenta dotted line demarcates proximal collective and distal dispersed NCLC. Note that upon XAV treatment all the NCLC are dispersed, whereas upon Chiron treatment all the NCLC display collective behaviour. (C) Schematic of the experiment (left). Passaged NCLC treated with Chiron or DMSO immunostained for β-catenin and TFAP2a. The graph represents cell density in individual colonies, n = 8 colonies from 3 biological experiments. Statistical analysis: Mann-Whitney test. *** is P=0.0006. Scale bar: 50μm. (D) NCLC-P2 treated with DMSO (top) or 3μM Chiron (bottom) immunostained to detect N-cadherin expression. Actin stained with Phalloidin to visualize cell morphology. (E) N-cadherin occupancy in cell membrane of NCLC treated with DMSO or Chiron. N-cadherin occupancy was calculated as the ratio of total length of N-cadherin staining to the sum of perimeter, of all the cells in the colony. Statistical analysis: Two tailed unpaired t test. * is P=0.0105. Scale bar: 100μm

Immunostaining of passaged NCLC revealed overall increase in β-catenin levels in Chiron treated cells. Notably, β-catenin was enriched at the cell-cell junctions. Also, TFAP2a expression was maintained upon Chiron treatment, suggesting retention of neural crest progenitor identity. However, in line with the increased β-catenin in cell-cell junctions, the cell density within colonies in the Chiron treated NCLC clusters was about 2 times higher compared to vehicle treated control (Figure 2C, n = 8 colonies from 3 biological replicates; *** is p<0.0005). Together, these observations suggest that β-catenin levels impact the collective versus dispersal behaviour of neural crest.

The correlation between increased β-catenin levels at the membrane and clustering of NCLC led us to investigate its role in regulating cell-cell junction. To do so, we measured the distribution of N-cadherin in the membrane of NCLC upon Chiron treatment. We observed that the tight cluster formation was associated with elevated N-cadherin occupancy on the membrane; two fold increase in the proportion of total membrane length occupied by N-cadherin staining upon treatment with Chiron (Figure 2 D,E ; n=10 and 12 colonies from 3 biological replicates for DMSO and Chiron treated conditions respectively; p=0.0105). This suggests increased formation of N-cadherin mediated cell-cell junction upon stabilisation of β-catenin levels. N-cadherin mediated stabilisation of cell-cell junction and clustering of NCLC is reminiscent of early collective behaviour of migratory CNCC in the developing embryo.

### Tissue-specific β-catenin stabilisation disrupts neural crest invasion of first pharyngeal arch

To test the causal relationship *in vivo* between modulation of β-catenin levels and collective / dispersed arrangement of CNCC, *w*e adopted a genetic approach. We used *Wnt1-Cre* and *Ctnnb1*^*lox(ex3)*^ alleles to constitutively stabilise β-catenin (*β-catenin*^*dEx3/+*^) specifically in neural crest. A reporter allele *ROSA*^*tdTomato*^ was included to mark the Cre-lox *‘*recombined cells’ (Fig. 3A, B). A previous study has reported that neural crest specific β-catenin gain of function leads to loss of distal skeletogenic CNCC, hypoplastic or partial formation of first pharyngeal arches and loss of all further caudal arches (Lee *et al.*, 2004). Notably, ectopic sensory neurogenesis was observed indicating that sustained Wnt / β-catenin signalling led to fate transformation of the presumptive distally-destined skeletogenic CNCC into sensory neurons (Lee *et al.*, 2004). Here, we revisited these mutants to specifically investigate the tissue organization of migratory and post-migratory CNCC. As previously reported, β-catenin gain-of-function mutant CNCC successfully undergo epithelial to mesenchymal transition and emigrate. However, while the proximal CNCC compartment at first pharyngeal arch level appeared slightly larger, the distal pool reaching the ventral arch location was drastically diminished (Fig. 3B, C). Strikingly, unlike the wild type embryos, N-cadherin levels appear uniform in the dorsoventral axis indicating a failure of cell dispersal in the distal end of migratory stream (Fig. 3C). In addition, no interstitial Fibronectin staining was observed in the diminished pool of distal mutant CNCC (Fig. 3D). Residual Tomato+ CNCC in the hypoplastic first pharyngeal arch expressed Twist1, a marker for skeletogenic or dermogenic CNCC (Goodnough, Dinuoscio and Atit, 2016; Fig. S6A, B). Presence of Twist1+ Tomato+ cells suggest that mutant CNCC may acquire the typical distal fate when they reach the arch environment. We also find that Twist1+ NCLC retain its expression upon Chiron-treatment (Fig. S6C). Based on these findings, we suggest that the previously reported fate transformation into ectopic sensory neurons in β-catenin gain-of-function could at least be in part due to mis-patterning; presumptive distal CNCC when mis-localised may fate transform in response to local cues. Taking this evidence together, we conclude that modulating membrane β-catenin levels is critical for appropriate CNCC patterning.

**Figure 3:**
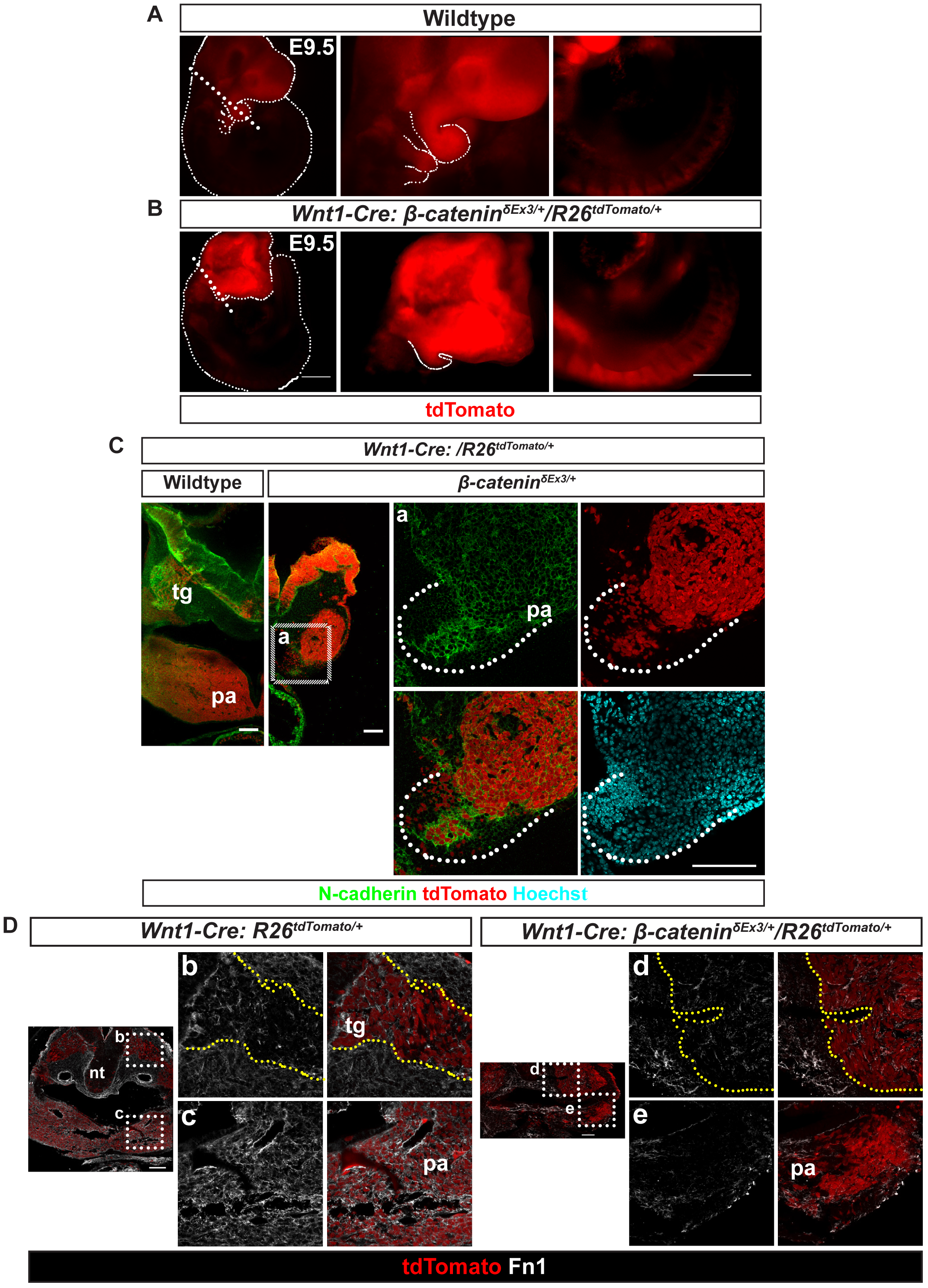
Tissue-specific β-catenin gain of function disrupts dispersal of cranial neural crest collective in mouse embryos. (A&B) Live reporter-tdTomato expression in *Wnt1-Cre:R26R*^*tdtTomato*^ embryos (A) or in *Wnt1-Cre:β-catenin*^*dex3/+*^;*R26R*^*tdtTomato*^ mutant embryos (B). Dotted lines mark the embryonic boundary (C) Transverse sections at the first pharyngeal arch axial level of E9.5 *Wnt1-Cre:R26R*^*tdtTomato*^ (right) or *Wnt1-Cre:β-catenin*^*δex3/+*^;*R26R*^*tdtTomato*^ (left) embryos immunostained to detect N-cadherin and tdTomato (live) expression. The higher magnification image of the region marked (a) is shown in the right. (D) Transverse sections at the level of first pharyngeal arches of E9.5 *Wnt1-Cre:R26R*^*tdtTomato*^ (left) or *Wnt1-Cre:β-catenin*^*δex3/+*^;*R26R*^*tdtTomato*^ (right) embryos immunostained to detect Fn1 and tdTomato (live) expression. Yellow dotted line in zoomed in images demarcate proximal CNCC domain. pa-pharyngeal arches, nt-neural tube, tg-anlage of trigeminal ganglia. Scale bar: 100μm

While our study highlights the function of β-catenin in the cell membrane, nuclear β-catenin levels were difficult to assess. Therefore, we assessed Wnt signal-driven nuclear β-catenin function using Wnt activity reporter mouse line *Axin2-d2eGFP.* While the reporter activity decreased in CNCC migrating distally at E8.5, no correlation of reporter expression was apparent with collective / dispersed states at E9.5 (Fig. S7A, B, C). Instead, *Axin2-d2eGFP* expression correlated with fate commitment within the proximal domain; expression of GFP and neural marker Tuj1 was mutually exclusive (Fig. S7D). Thus, the collective / dispersed architecture and the varying membrane β-catenin levels, do not appear to correlate with nuclear β-catenin function. Moreover, the results also uncover the lack of correlation between Wnt reporter expression and membrane β-catenin levels. In fact, Wnt signal independent function of β-catenin is well documented in various developmental context (Haq *et al.*, 2003; Ding *et al.*, 2005). Taken together, the correlative evidence from the mouse embryos and the causal evidence provided by the functional studies, both in mouse embryos and human NCLC model, underscore a novel role of β-catenin in regulating the reorganization of CNCC architecture by modulating cell-cell adhesion (Fig. 4), which is intricately linked to morphogenesis of CNCC derived structures.

**Figure 4.**
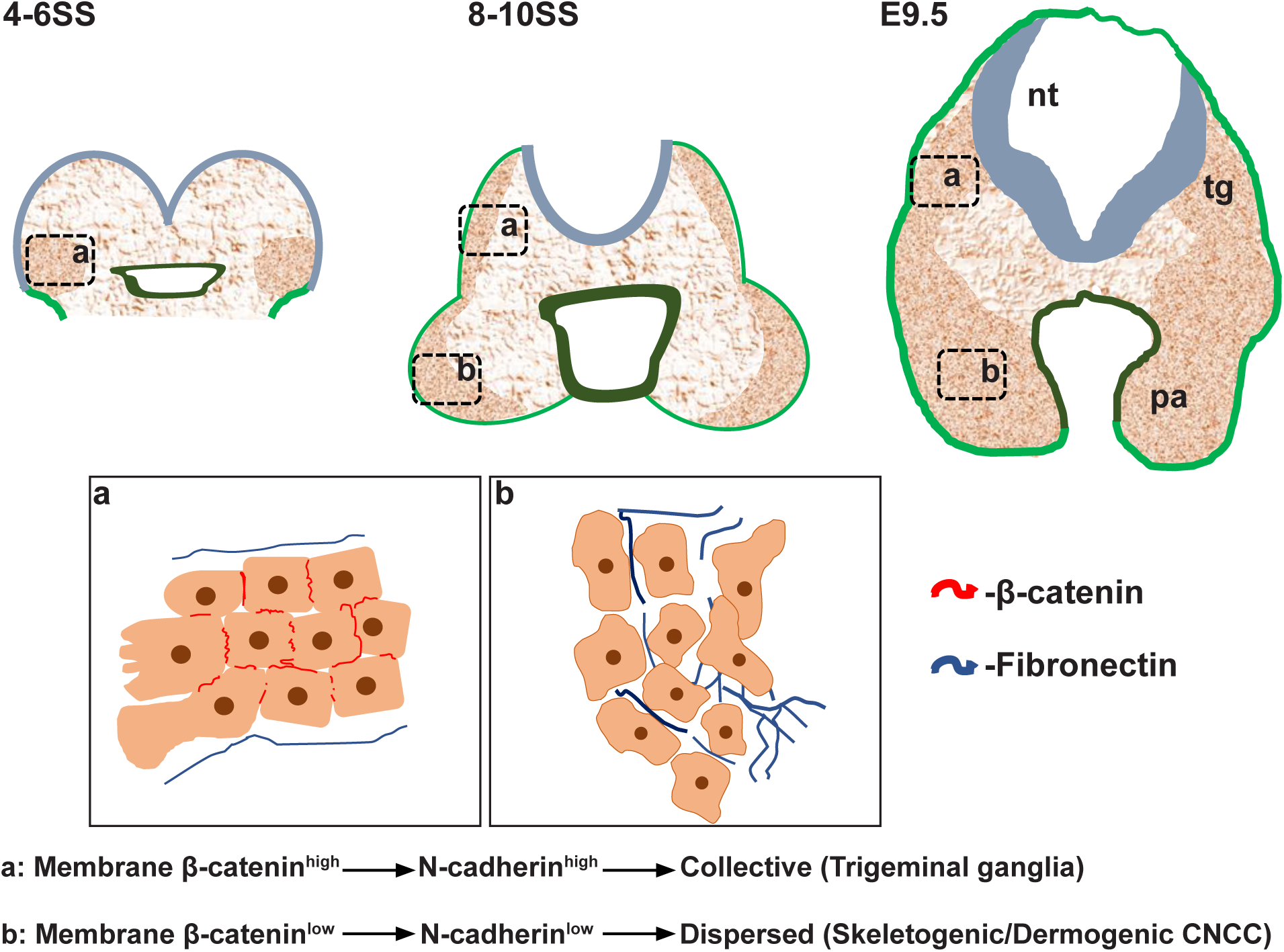
Modulation of β-catenin levels is critical for patterning of CNCC: A schematic describing the proposed model. As the CNCC emerge from the neural plate border, cells retain cell-cell contact to display a collective behaviour. With the progression of development, proximal CNCC retain collective behaviour whereas the distal CNCC disperse into the pharyngeal arches. The switch is regulated by membrane β-catenin levels via N-cadherin mediated adherens junctions. pa-pharyngeal arches, nt-neural tube, tg-anlage of trigeminal ganglia.

Canonical Wnt/β-catenin signalling is a major signalling pathway that regulates CNCC development at multiple levels. Earlier studies have demonstrated the role of Wnt/β-catenin signalling in CNCC induction, delamination, fate choice and also terminal differentiation of downstream lineages (Martin I. García-Castro, Christophe Marcelle, 2002; Lee *et al.*, 2004). In this study using mouse genetics approach and human *in vitro* cell culture system, we have demonstrated a novel regulatory role of β-catenin in tissue level organization of CNCC.

Transitions in tissue organization from epithelial to mesenchymal state or vice versa is known to be a major driver of morphogenesis during embryonic development (Thiery, 2003). A recent study has reported that transitions within mesenchyme between collective and dispersed state is associated with solid-like to fluid-like tissue characteristic, respectively. Moreover, this change in physical property has been demonstrated to drive tissue elongation, thus, enabling axial extension during posterior body morphogenesis in zebrafish (Mongera *et al.*, 2018). Such transitions are proposed to be more widespread during embryonic development. Here, we report a similar transition in tissue architecture in the migratory stream of CNCC. Our observations suggest that, at the site of origin, subsequent to epithelial to mesenchymal transition, CNCC retain cell-cell contact. In fact, such a collective state is important for directional migration of CNCC as postulated by two competing computational models (Kulesa *et al.*, 2010; Szabó and Mayor, 2016). We provide evidence for the collective arrangement in mice and human and show that, as development progresses the CNCC collective disperses distally. We also reveal the molecular basis for the tissue architecture transition; however, the signal regulating β-catenin levels is yet to be addressed.

Failure to disperse in mutants with neural crest specific stabilization of β-catenin appear to underlie their failure to invade the 1^st^ pharyngeal arch. Diminished number of CNCC arriving ventrally at the arches as well as the possible loss of signals from these cells may explain the hypoplastic 1^st^ arch and the failure to form caudal arches. In summary, our results reveal a patterning event during CNCC development, which involves compartment-specific transition of CNCC from collective to dispersed state. The process regulated through modulation of β-catenin levels appears to control CNCC influx into pharyngeal arches. These findings uncover a novel mechanism governing CNCC patterning.

## Acknowledgements

We thank animal facility (ACRC-NCBS/inStem), Central Imaging & Flow Cytometry Facility (NCBS) and Stem Cell Facility (inStem). We thank Profs. Makoto M Taketo and Shubha Tole for the Catnb^lox(ex3)^, Prof. Frank Costantini for Axin2-d2eGFP mouse line. We thank Drs. Vanitha R and Jayaprakash P for technical help. We thank Profs. Philippa Francis-West and Amitabha Bandhyopadhyay for discussion.

## Author contributions

Conceptualization: A.J., R.S.; Methodology: A.J., V.L.; Validation: A.J., V.L.,; Formal analysis: A.J., V.L., D.P., R.S.; Investigation: A.J., V.L.; Writing -original draft: A.J., R.S.; Writing -review & editing: A.J., R.S.; Visualization: A.J., V.L., R.S.; Supervision: R.S.; Project administration: R.S.; Funding acquisition: R.S.

## Data availability

Transcriptome data of H9-ESC, Neurospheres and NCLC will be deposited in the Sequence read archive (SRA).

## Materials and Methods

### Animal handling

All experiments on mice were carried out in compliance with the Committee for the Purpose of Control and Supervision of Experiments on Animals (CPCSEA) national guidelines. All experimental procedures were approved by the InStem Institutional Animal Ethics Committee.

### Cell culture

#### Human ES cell maintenance

H9 cells (WiCell, SLA# 16-W0026) were cultured in mTESR1 (Stem Cell Technologies). Cultured cells were replenished with fresh media every day. Passaging of the cells was performed at 1:3 ratio when the cells attained approximately 70-80% confluency. For passaging, 70-80% confluent cultures were treated with CTK till the edges of the colonies detached. CTK was then aspirated out, the enzymatic reaction was neutralized by mTESR1, colonies were gently scratched, collected and were spun down by short centrifugation of 10 seconds. Cells were then resuspended in fresh mTESR and seeded on the Matrigel coated plates.*

For differentiation, we adapted a published protocol in Bajpai *et al.*, 2010. We started with 70-80% confluent H9 ESCs for NCLC differentiation. CTK treatment followed by subsequent steps till the collection of H9 colonies was performed as above. Collected cell colonies were resuspended at 1:1 ratio in neural induction media supplemented with growth factors and were grown in suspension using low attachment plates. Media was changed every day for first five days during which majority of the colonies organized into spherical aggregates called neurospheres. Subsequently, media was changed on alternate days. Neurospheres were maintained in suspension culture until the appearance of rosette-like structures within all the spheres. Once the majority of neurospheres began to form rosettes, typically in 9 – 11 days of suspension, they were seeded on the culture plates coated with Fibronectin (1 µg / ml). Upon attachment, migratory NCLC begin to emerge from neurospheres within 12 hrs. These are termed as P0-NCLC.

To passage migratory NCLC emerging out from attached neurospheres, syringe needles were used to mechanically separate migratory cells from neurospheres and detach them from the plates. Media, along with the detached spheres were aspirated out. The attached cells were treated with CTK for 10 seconds (1 ml CTK per 60 mm culture dish). CTK was then aspirated out. Cells were then collected in fresh media by gentle pipetting (2 ml media per 60 mm dish). Cells suspended in the media was then collected in 15 ml conical tubes and the cells were spun down for 3 minutes at 1000 RPM. Spent media was aspirated out, cells were then resuspended in appropriate amount of fresh media (NIM+GF) and seeded on fibronectin-coated dishes. First passaging was done at 1:1 ratio. Significant cell death is observed during this step of NCLC culture, potentially due to residual neurospheres left over on the plate after mechanical removal, which fail to attach after passaging. Initially, media was changed 24 hours after passaging and subsequently, media was changed on alternative days. For subsequent passaging of NCLC, around 80% confluent cultures were treated with CTK for 30 seconds, rest of the steps were as described for the first passaging of NCLC. Seeding of NCLC on fibronectin is done at 1:3 ratio on the fibronectin-coated plates.** All the experiments involving passaged NCLC were performed with P2 or P3 NCLC to avoid heterogeneity in the culture as the culture becomes uniformly Twist1 positive beginning from second passage. After first passage ∼20% of the NCLC express Sox10 (Figure S8).

*Matrigel dilutions were made as described in the certificate of analysis of each batch. For coating Matrigel, aliquots were diluted in DMEM (as recommended by the manufacturers), and spread evenly throughout the plates. The plates were then incubated at 37°C for at least two hours. Matrigel solution can be replaced with fresh media and plates can be sealed and stored in 4°C for up to one week.

**Fibronectin (Sigma, F1141), was diluted in PBS at a concentration of 1 μg / ml concentration, and coating was done as described previously for Matrigel coating. Fibronectin solution can be replaced with PBS and plates can be stored in 4°C for up to one week.

### Immunostaining

#### Embedding and sectioning of embryos

Mouse embryos were dissected in 1X PBS. For E8.5 or younger embryos, 1X PBS + 0.1%Tween 20 (1X PTW) was used to prevent embryos sticking to each other. The dissected embryos were fixed in 4% PFA at 4°C for 90 minutes. All traces of PFA was then removed by three rinses and three washes of ten minutes each using 1X PTW. Embryos were then equilibrated with 15% sucrose prepared in PBS by incubating overnight at 4°C. The equilibrated embryos were transferred to a tube containing 7.5% high bloom gelatin solution (300 bloom SuperClear™ Gelatin obtained from Custom Collagen, USA) prepared in 15% Sucrose. The tubes were incubated at 37°C at least for one hour. Individual embryos were then transferred to each mould and then appropriately oriented. The blocks were allowed to solidify at 4°C. The solid blocks were cryo-frozen by inserting the blocks in the liquid nitrogen for about 15 seconds. The frozen blocks were either cryo-sectioned immediately or stored in -80°C freezer in air-tight container. The embryos were sectioned at 16 µm thickness using a Leica CM 1850 UV cryostat. The sections were collected on the charged slides (Leica SuperFrost plus, Ref: 6776214).

#### Immunostaining

Sections were blocked and permeabilized for at least an hour using blocking solution: with 10% normal donkey serum in PBS containing 0.5% Triton X-The embryos were then treated overnight at 4°C with primary antibodies (see Table 5) diluted in the blocking solution, followed by 1 h incubation in PBS containing 0.1% Tween 20, 10 mg / ml Hoechst and 4 μg / ml secondary antibodies. For Phalloidin staining, Alexa Fluor™ 488 Phalloidin (Life technologies, A12379; 1:100) was added to the solutions containing secondary antibodies.

For immunostaining cultured cells, the cells were fixed with 4% PFA for 15 min at ambient temperature and permeabilized using 0.5% Triton X-100 in PBS for 30 min and further blocked in PBS with 0.5% Triton X-100 and 10% normal donkey serum for 1 h. Incubation with primary antibody in blocking solution was carried out overnight at 4°C. The secondary antibody treatment was performed as described for immunostaining of sections.

#### Image acquisition and analysis

Immunostained cryosections were imaged using Olympus FV1000 or FV3000 confocal microscope. All image analyses were performed using Fiji (Schindelin *et al.*, 2012). To measure the mean grey value of β-catenin and N-cadherin in membrane, free hand selection tool was used to mark the outer and inner surface of the membrane of neural crest cells determined by the expression of tdTomato reporter. Mean grey value was calculated using the following formula:

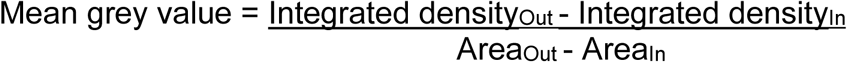

To measure the cell density of NCLC, overall area occupied by a CNCC colony was measured by drawing a boundary by free hand tool around a colony. The number of nuclei were counted manually using Hoechst channel. The number of cells were divided by colony area to calculate the density of the cells within the colony.

To measure N-cadherin occupancy, images were manually thresholded to specifically mark N-cadherin expression in the cell membrane. The number of pixels corresponding to N-cadherin expression in membrane were measured. This was divided by the sum of perimeter of all the cells, measured by drawing a boundary around individual cells using free-hand tool in a Phalloidin channel of the same image, to get N-cadherin occupancy.

We measured the mean GFP intensity in migratory CNCC in transverse sections of E8.5 *Axin2-d2eGFP*, by marking boundaries around the TFAP2a positive cells. The distance between the lateral posterior tip of the section and the centroid of each cell was measured as a proxy for distance of cells from the site of origin.

### Transcriptome analysis

Total RNA (1 µg) extracted from NCLC and neurospheres, using the Trizol reagent was used to prepare library for sequencing. The library was quality checked using Quibit and Bioanalyzer. After passing QC, we performed paired-end RNA sequencing (2 × 75 bps) for NCLC and neurospheres. We obtained ∼27 to 45 million reads per sample. Adapters were trimmed using Trimmomatic (Bolger, Lohse and Usadel, 2014) and subsequently mapped to rRNA. Reads that did not align to rRNA were taken for further analysis. Reference-based transcriptome assembly algorithms Hisat2 v2.1.0; Cufflinks v2.2.1 and Cuffdiff v2.2.1 (Trapnell *et al.*, 2010, 2012; Kim, Langmead and Salzberg, 2015) was used to identify differentially expressed transcripts. We aligned the reads to human (hg38) genome-using Hisat2 (-q-p8--min-intronlen 50--max-intronlen 250000--dta-cufflinks--new-summary—summary file).

Around 96-97% of reads mapped to reference genome. The mapped reads were assembled using Cufflinks using hg38 Refseq gtf file. Transcripts which had adjusted *P*-values < 0.05 and minimum two-fold up/down regulated were considered as significantly expressed and taken for further analysis. We performed pathway analysis and gene-ontology analysis for these selected up/down regulated transcripts using gene set enrichment analysis (GSEA) (Subramanian *et al.*, 2005; Mi *et al.*, 2017). We used customized perl script for all the analysis used in this study. We used R ggplot2 (Wickham, 2010) & CummeRbund library for plotting.

### qPCR

Total RNA (500 ng-1 µg) extracted using the Trizol (Invitrogen) was used to prepare cDNA (Super-Script III, Invitrogen). Oligo dT primers were used for cDNA synthesis from total RNA. Real-time PCR reactions were performed using PowerSYBR Green Universal Master Mix (Thermo Scientific) and analysed using ViiA7 Real Time PCR system (Applied Biosystems). Data were normalized using GAPDH expression. The specificity of the primers was tested by melt curve. The primers are listed in Table 6. Each condition was tested using two technical replicates and at least three biological replicates was analysed for relative gene expression using 2-ΔΔCt method (Schmittgen and Livak, 2008).

