## Supplementary information for "Modulation of β-catenin levels is critical for cranial neural crest patterning and dispersal into first pharyngeal arch"

**Supplementary figures:**

#### Figure S1

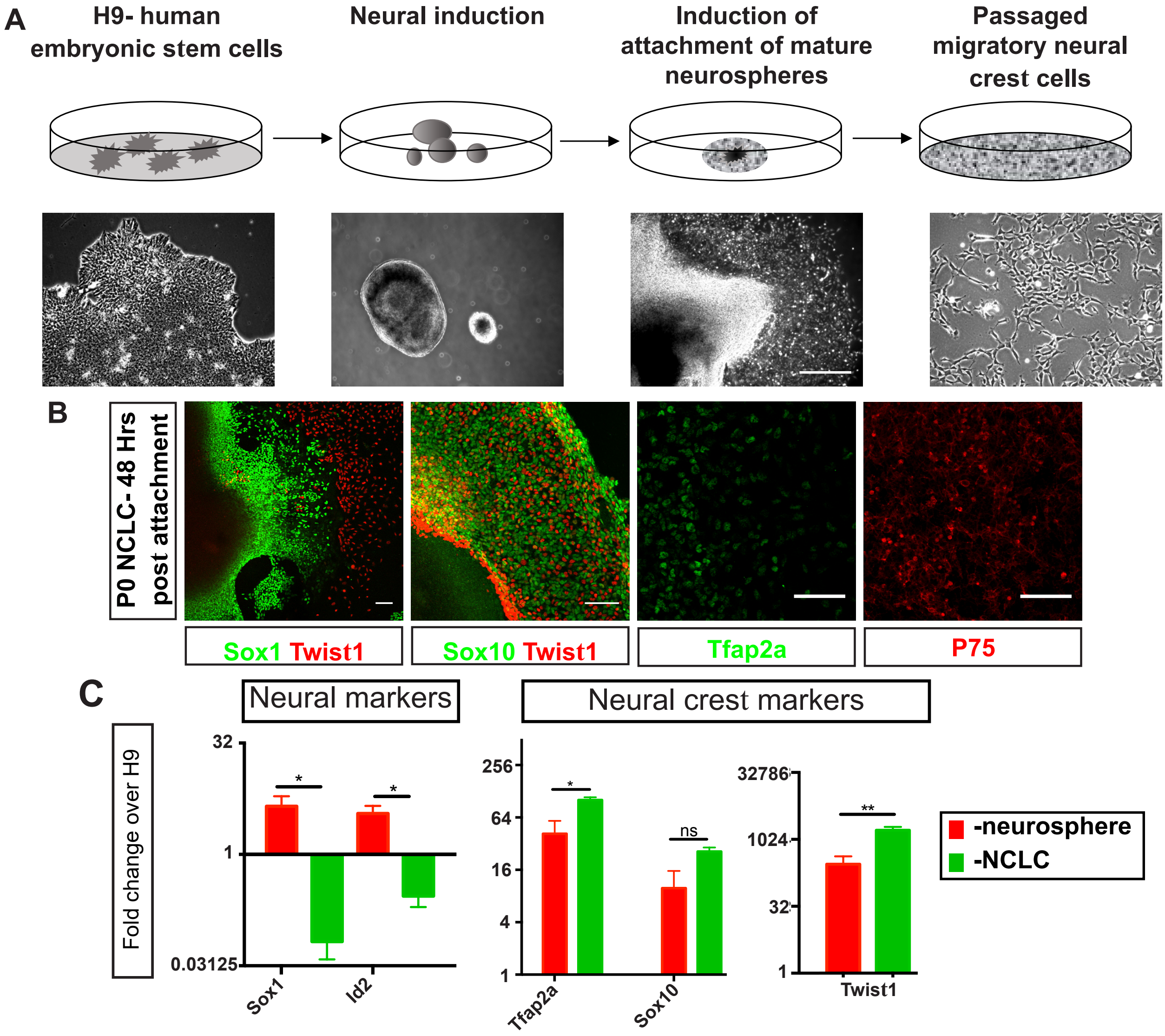

A
H9- human embryonic stem cells
Neural induction
Induction of attachment of mature neurospheres
Passaged migratory neural crest cells
B
P0 NCLC- 48 Hrs post attachment
Sox1 Twist1
Sox10 Twist1
Tfap2a
P75
C
Neural markers
Neural crest markers
Fold change over H9
32
1
0.03125
Sox1
Id2
256
64
16
4
1
Tfap2a
Sox10
ns
32786
1024
32
1
Twist1
-neurosphere
-NCLC

#### Figure S2

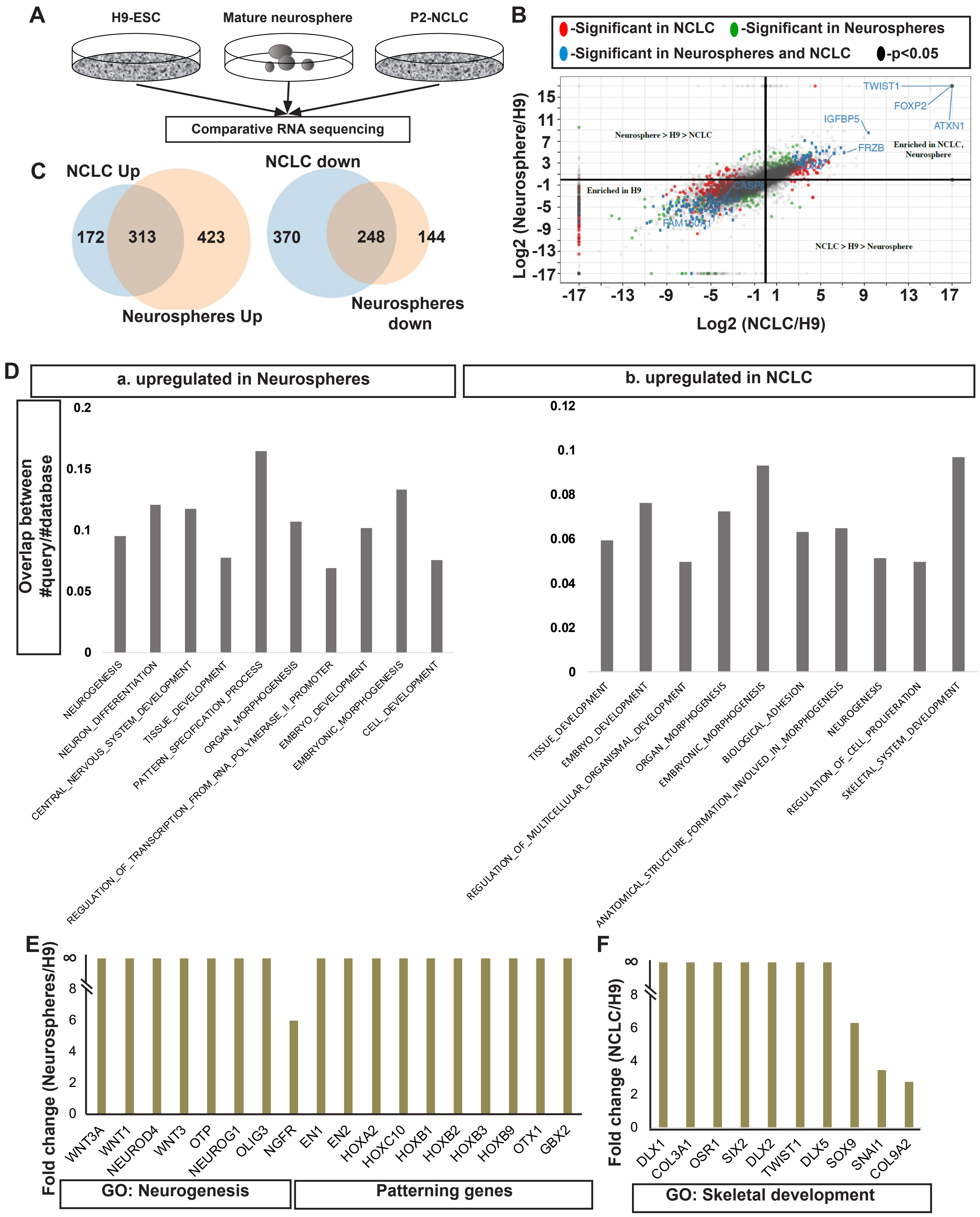
A
H9-ESC
Mature neurosphere
P2-NCLC
Comparative RNA sequencing
B
-Significant in NCLC
-Significant in Neurospheres
-Significant in Neurospheres and NCLC
-p<0.05
Log2 (Neurosphere/H9)
Log2 (NCLC/H9)
Neurosphere > H9 > NCLC
Enriched in H9
Enriched in NCLC, Neurosphere
NCLC > H9 > Neurosphere
TWIST1
FOXP2
ATXN1
IGFBP5
FRZB
CASP8
FAM150A1
C
NCLC Up
172
313
423
Neurospheres Up
NCLC down
370
248
144
Neurospheres down
D
a. upregulated in Neurospheres
b. upregulated in NCLC
Overlap between #query/#database
NEUROGENESIS
NEURON_DIFFERENTIATION
CENTRAL_NERVOUS_SYSTEM_DEVELOPMENT
TISSUE_DEVELOPMENT
PATTERN_SPECIFICATION_PROCESS
ORGAN_MORPHOGENESIS
REGULATION_OF_TRANSCRIPTION_FROM_RNA_POLYMERASE_II_PROMOTER
EMBRYO_DEVELOPMENT
EMBRYONIC_MORPHOGENESIS
CELL_DEVELOPMENT
TISSUE_DEVELOPMENT
EMBRYO_DEVELOPMENT
REGULATION_OF_MULTICELLULAR_ORGANISMAL_DEVELOPMENT
ORGAN_MORPHOGENESIS
EMBRYONIC_MORPHOGENESIS
BIOLOGICAL_ADHESION
ANATOMICAL_STRUCTURE_FORMATION_INVOLVED_IN_MORPHOGENESIS
NEUROGENESIS
REGULATION_OF_CELL_PROLIFERATION
SKELETAL_SYSTEM_DEVELOPMENT
E
Fold change (Neurospheres/H9)
WNT3A
WNT1
NEUROD4
WNT3
OTP
NEUROG1
OLIG3
NGFR
EN1
EN2
HOXA2
HOXC10
HOXB1
HOXB2
HOXB3
HOXB9
OTX1
GBX2
GO: Neurogenesis
Patterning genes
F
Fold change (NCLC/H9)
DLX1
COL3A1
OSR1
SIX2
DLX2
TWIST1
DLX5
SOX9
SNAI1
COL9A2
GO: Skeletal development

### Figure S3

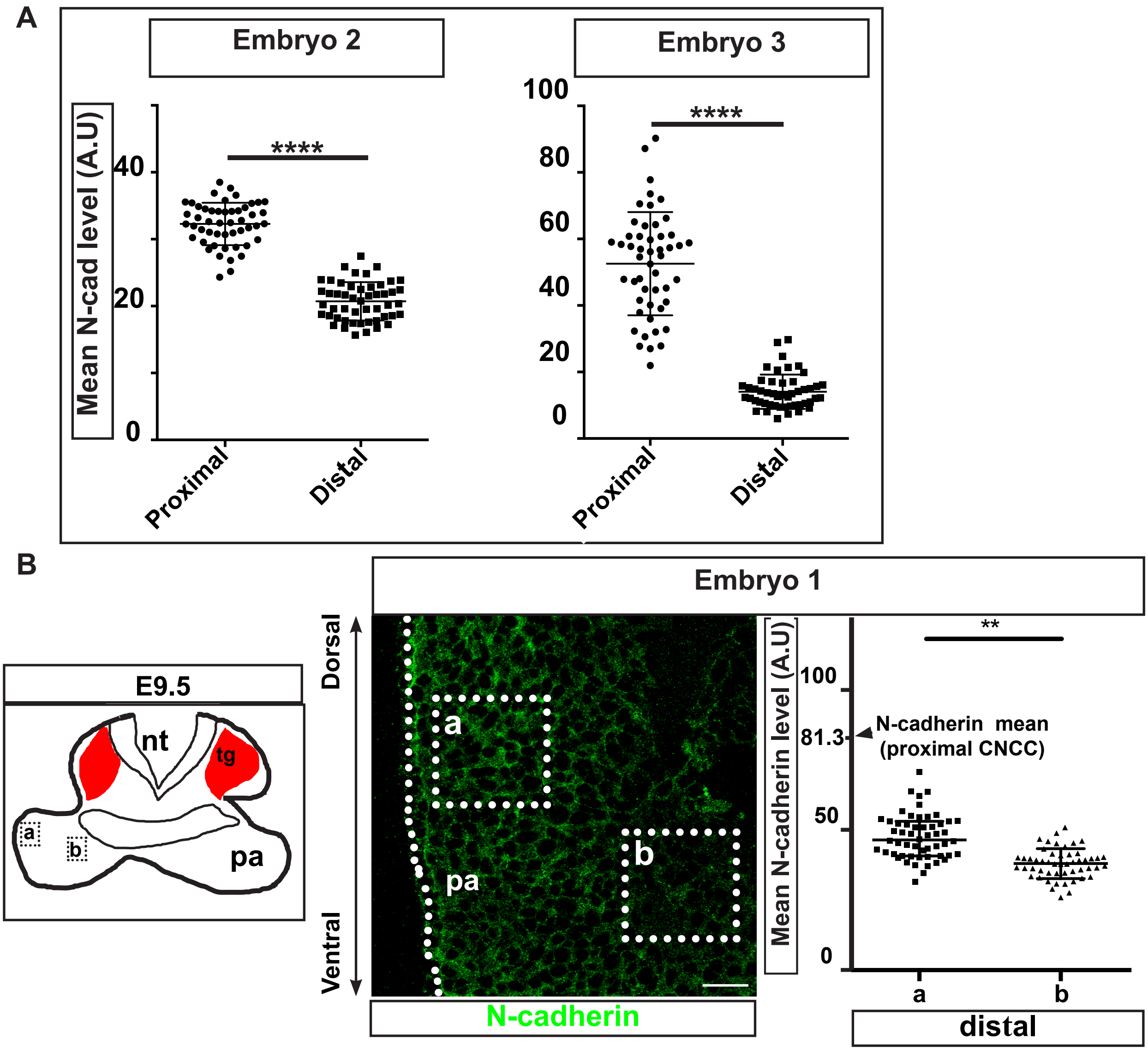

A
Embryo 2
Embryo 3
Mean N-cad level (A.U)
40
20
0
****
100
80
60
40
20
0
****
Proximal
Distal
Proximal
Distal
B
E9.5
nt
tg
a
b
pa
Embryo 1
Dorsal
Ventral
a
b
pa
N-cadherin
Mean N-cadherin level (A.U)
100
81.3
50
0
N-cadherin mean
(proximal CNCC)
**
a
b
distal

### Figure S4

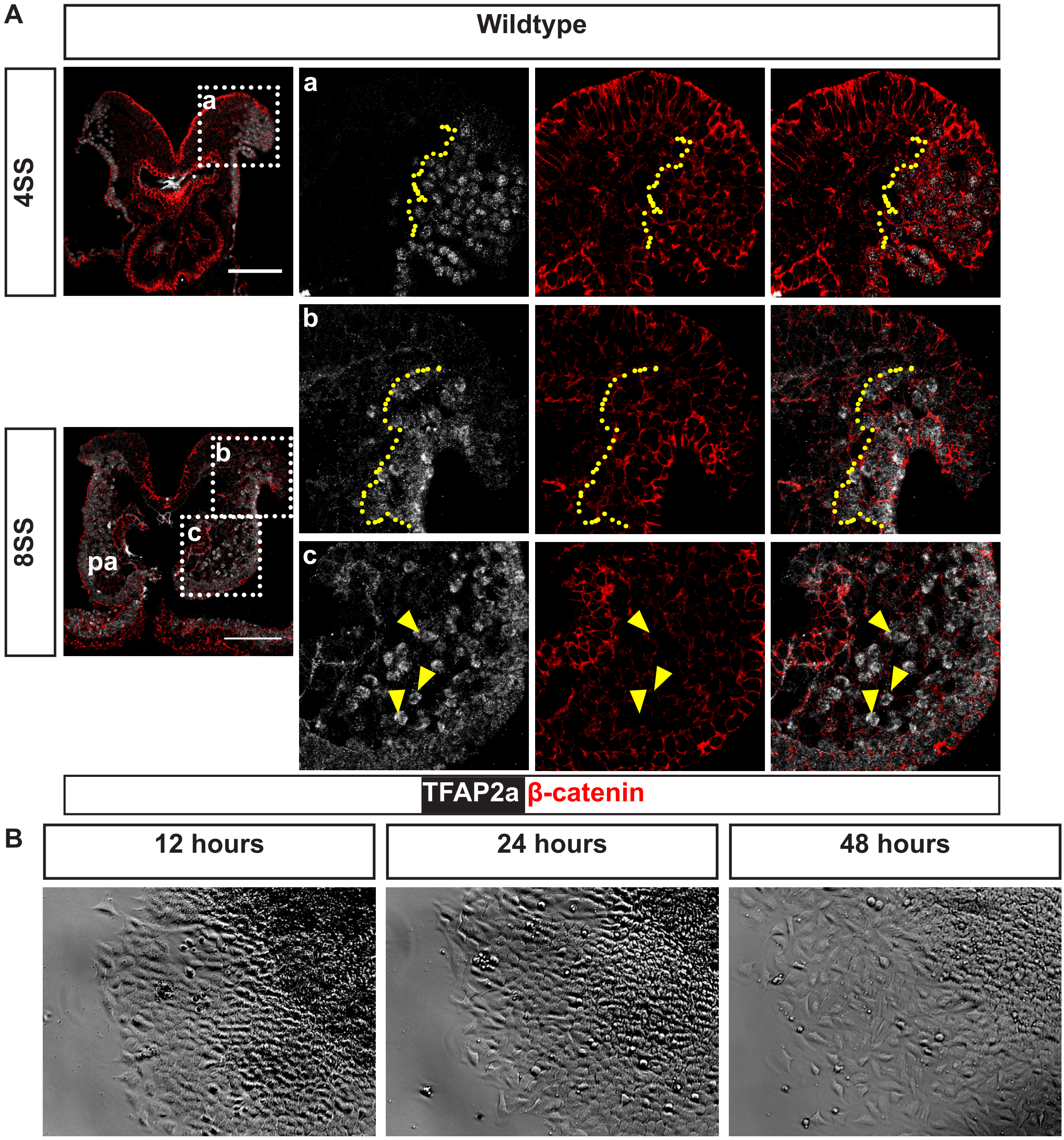
A
Wildtype
4SS
8SS
a
b
c
pa
TFAP2a β-catenin
B
12 hours
24 hours
48 hours

#### Figure S5

A

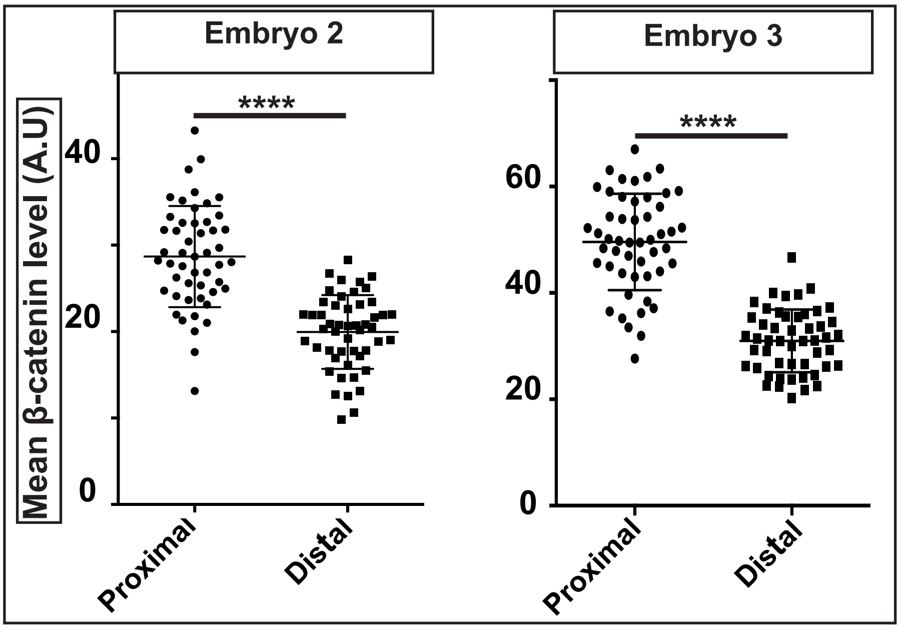

Embryo 2
Embryo 3
Mean β-catenin level (A.U)
****
****
40
20
0
60
40
20
0
Proximal
Distal
Proximal
Distal

#### Figure S6

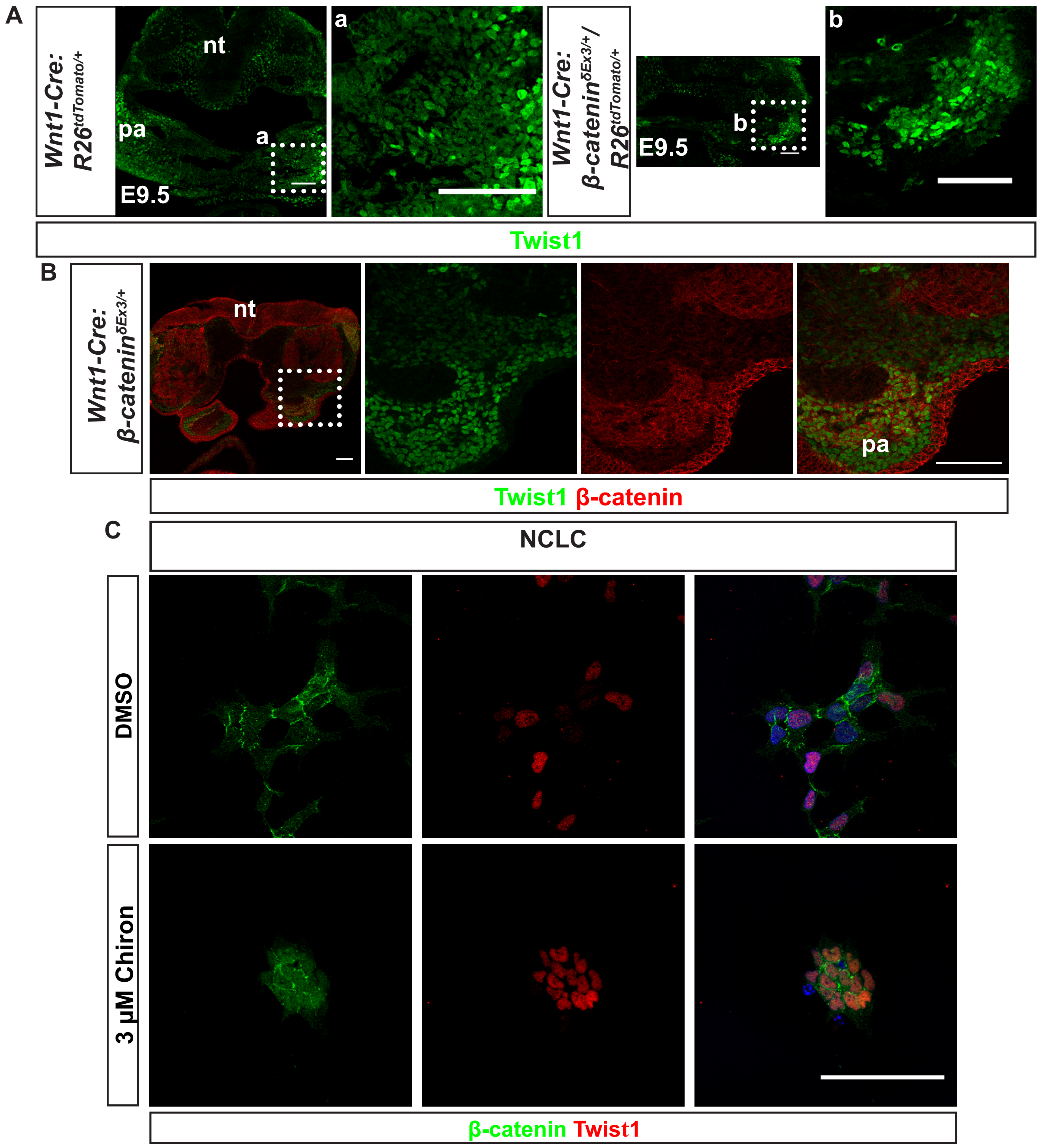

A
Wnt1-Cre: R26tdTomato/+
nt
pa
a
E9.5
a
Wnt1-Cre: β-cateninδEx3/+/ R26tdTomato/+
E9.5
b
b
Twist1
B
Wnt1-Cre: β-cateninδEx3/+
nt
pa
Twist1 β-catenin
C
NCLC
DMSO
3 µM Chiron
β-catenin Twist1

### Figure S7

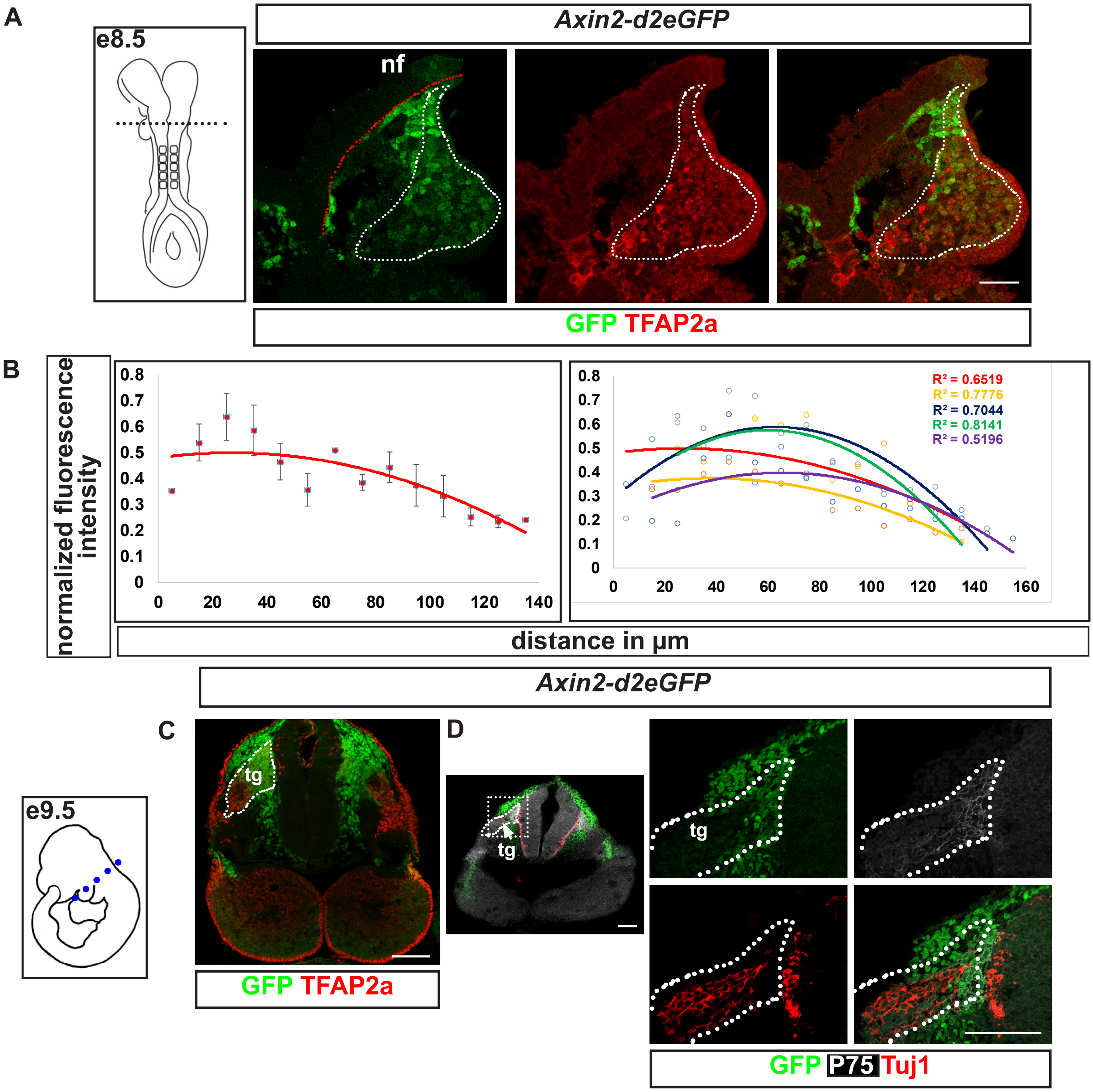
A
e8.5
Axin2-d2eGFP
nf
GFP TFAP2a
B
normalized fluorescence intensity
0.8
0.7
0.6
0.5
0.4
0.3
0.2
0.1
0
0 20 40 60 80 100 120 140
0 20 40 60 80 100 120 140 160
R² = 0.6519
R² = 0.7776
R² = 0.7044
R² = 0.8141
R² = 0.5196
distance in µm
Axin2-d2eGFP
C
e9.5
tg
GFP TFAP2a
D
tg
tg
GFP P75 Tuj1

Figure S8

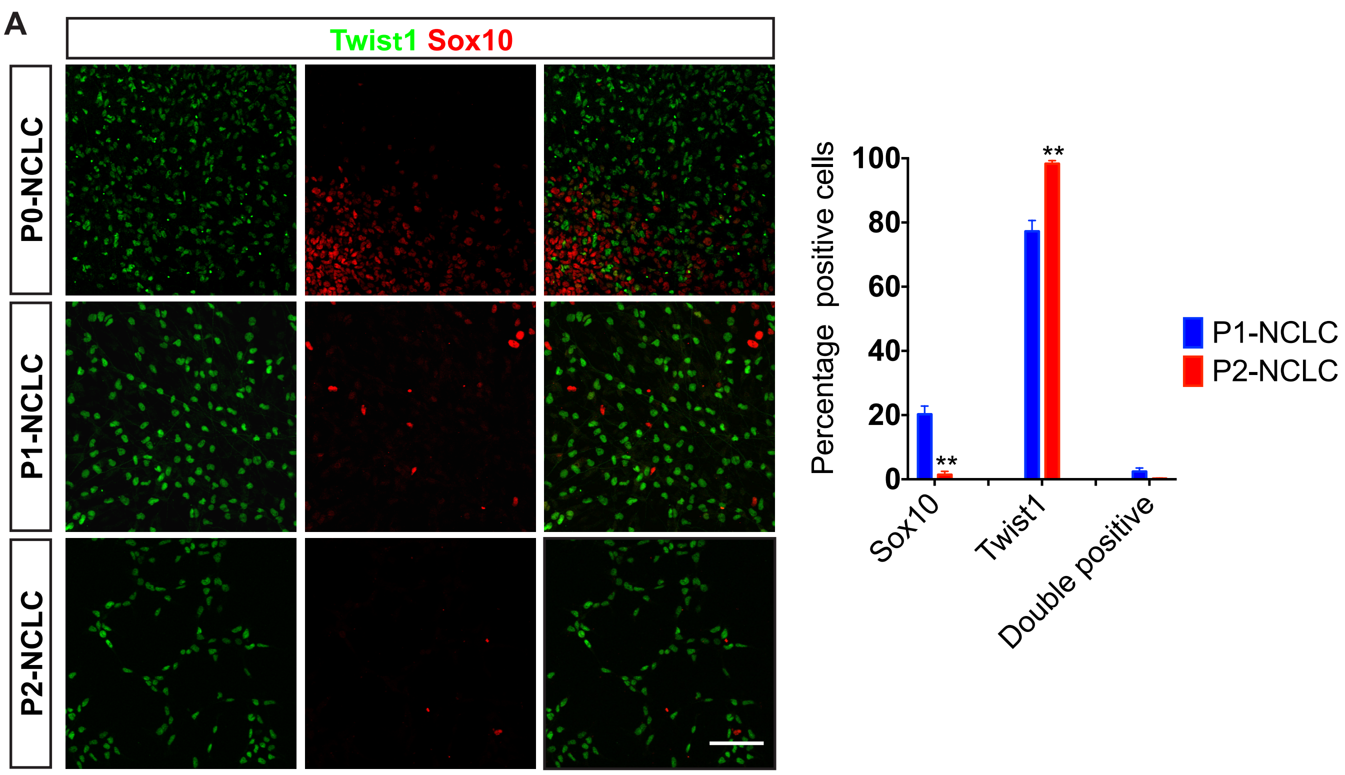
A
Twist1 Sox10
P0-NCLC
P1-NCLC
P2-NCLC
Percentage positive cells
100
80
60
40
20
0
**
**
Sox10
Twist1
Double positive
P1-NCLC
P2-NCLC

**Supplementary figure legends:**

**Figure S1. Human embryonic stem cell derived neural crest-like cells provide amenable model to study neural crest development.** (A) Schematic of differentiation of human embryonic stem cells into NCLC (top) and bright field images of colonies of corresponding stages of differentiation. (B) Migratory NCLC emerging from attached neurospheres immunostained to detect various neural and neural crest markers. Note that migratory NCLC uniquely express neural crest markers such as Twist1/Sox10, TFAP2a and p75 and downregulate neural markers Sox1 and Sox2. (C) Real time quantitative PCR analysis from H9- ESC, neurospheres before attachment and passage two NCLC shows the dynamics of neural and neural crest marker expression. Statistical analysis: one-way ANOVA with multiple comparisions, * $p<0.05$, ** $p<0.005$, error bars indicate S.E.M. Scale bar: 100μm

**Figure S2. Comparative RNA-seq analysis globally validates NCLC as CNCC model.** (A) Schematic showing the sample used for comparative RNA sequencing analysis between mature neurospheres, P2-NCLC and H9-ESC. (B) RNA-seq analysis was performed in biological duplicate samples. The scatter plot shows the distribution of differentially expressed genes. (C) Venn diagram showing number of differentially expressed genes upregulated (left), downregulated (right) between NCLC and H9 or neurospheres and H9. (D) Gene set enrichment analysis of differentially expressed genes between neurospheres and H9 (a) or NCLC and H9 (b). Y-axis is the overlap between the differentially expressed genes from the transcriptome analysis and gene-set associated with a specific GO term in the database. It is measured as the proportion between number of differentially expressed genes present in a gene-set associated with a specific GO term to the total number of genes present in a gene-set associated with that GO term in the database. (E&F) Fold change of selected transcripts from RNA seq data set. The comparison is between neurosphere vs H9 (E) or NCLC vs H9 (F).

**Figure S3. Differential N-cadherin expression reveals distinct cell arrangements in fate specific CNCC compartments.** (A) Quantitation of N-cadherin expression in the membranes of CNCC in the proximal and distal regions at the level of first pharyngeal arches in E9.5 mouse embryos in embryos 2 and 3. (B) Difference in N-cadherin levels within the 1$^{st}$ pharyngeal arch. Pharyngeal arch region zoomed from

N-cadherin immunostained section of embryo 1, from Figure 1C. Quantitation of N-cadherin levels in the membrane from the regions a and b (marked in the schematic) shows differential expression in CNCC within pharyngeal arch. Closer to epidermis, CNCC express higher level of N-cadherin relative to the cells dispersed away from the epidermis. Fibronectin distribution within arch appears to be in accordance with that of N-cadherin. Dotted line marks the lateral epidermis. pa- pharyngeal arch, nt- neural tube. Statistical analysis: two tailed unpaired t test. **** is p<0.00001.** is p<0.001.

**Figure S4. Migratory CNCC undergoes transition from collective to dispersed state in mouse embryos.** This behaviour is recapitulated in human NCLC *in vitro.* (A) Time series analysis of migratory CNCC at the level of prospective first pharyngeal arch. Transverse sections of 4 somite stage (4SS) and 8 somite stage (8SS) mouse embryos were immunostained for β-catenin and TFAP2a. Yellow dotted line demarcates CNCC collective. Yellow arrowhead marks distally dispersed CNCC. PA – pharyngeal arch. (B) Bright field images of migratory NCLC emerging from attached neurospheres at 12, 24 and 48 hours. Note that at 12 hours NCLC maintain cell-cell contact. Distally located cells begin to disperse at 24 hours. pa- pharyngeal arch. Scale bar: 100μm.

**Figure S5. Reduction in β-catenin levels in cell membrane correlates with reduction in N-cadherin levels.** Companion Figure to Figure 1F showing data from the replicates. (A) Quantitation of membrane β-catenin levels in the proximal and distal CNCC at the level of first pharyngeal arches in E9.5 mouse embryos 2 and 3. Note the correlation between the levels of β-catenin and N-cadherin (Refer Supplementary Figure 3A). Statistical analysis: two tailed unpaired t test. **** is p<0.00001.

**Figure S6. Stabilisation of β-catenin is not sufficient for fate transformation in CNCC lineage.** (A) Transverse sections at the level of first pharyngeal arches of E9.5 *Wnt1-Cre:R26R$^{tdtTomato}$* or *Wnt1-Cre:β-catenin$^{\delta ex3/+}$;R26R$^{tdtTomato}$* embryos immunostained to detect Twist1. Same sections as shown in Figure 3D; see 3D for reporter expression. Twist1 is expressed in cranial mesoderm as well; reporter expression helps assess expression in neural crest. (B) Transverse sections at the level of first pharyngeal arches of E9.5 *Wnt1-Cre:β-catenin$^{\delta ex3/+}$* embryo immunostained to detect Twist1. Note Twist1+ cells show high β-catenin levels in the hypoplastic pharyngeal arch. (C) NCLC treated with DMSO (top) or 3μM Chiron (bottom) immunostained to detect β-catenin and Twist1. pa- pharyngeal arch, nt- neural tube. Scale bar: 100μm.

**Figure S7. Wnt activity reporter mouse indicates differential dynamics of nuclear and membrane β-catenin.** (A) Immunostaining to detect GFP and TFAP2a expression in transverse section of E8.5 *Axin2-d2eGFP* mouse embryo at the level of first pharyngeal arch. White dotted line marks the spread of migratory neural crest cells in the section, red dotted line demarcates the neural fold (nf). (B) Plots show normalized GFP intensity in individual TFAP2a+ neural crest cells. Each point in the left plot shows the mean of normalized GFP intensity value of the cells binned as per their position relative to the dorsal edge of neural fold; error bars represent standard deviation. Right plot shows the trend lines measured from five images, from three different embryos. (C&D) Immunostaining to detect GFP and TFAP2a (C) or GFP, P75 and Tuj1 (D) expression in transverse section of E9.5 stage *Axin2-d2eGFP* mouse embryo at the level of first pharyngeal arch. pa- pharyngeal arch, tg- anlage of trigeminal ganglia, nf- neural fold. Scale bar: 100μm.

**Figure S8. Homogenous Twist1 expression in passaged NCLC.** (A) Immunostaining of migratory NCLC at different passages to detect expression of Twist1 and Sox10. (B) Quantitation of the number of cells expressing either of Twist1 or Sox10 or both in the culture (n = 3 independent experiments). Note, P2-NCLC form homogenous Twist1 expressing culture. Statistical analysis: non-parametric unpaired t-test, ** is $p<0.005$, **** is $p<0.00005$, error bars S.E.M. Scale bar: 100 μm.

**Table 1: Animal strains.**

| Strain | Reference | Source / Provider |
|---|---|---|
| *Wnt1-Cre* | Lewis *et al.*, 2013 | The Jackson Laboratory, USA, Stock # 022137 |
| *Rosa-tdTomato* | Madisen *et al.*, 2010 | The Jackson Laboratory, USA, Stock # 007914 |
| *β-catenin*$^{loxEx3}$ | Harada *et al.*, 1999 | Makoto Mark Taketo, Kyoto University, Japan |
| *Axin2-d2eGFP* | Jho *et al.*, 2002 | Frank Costantini, Columbia University Irving Medical Center, USA |

**Table 2: Composition of CTK solution.**

| Component | Final Concentration | Source | Catalogue number |
|---|---|---|---|
| Collagenase-IV | 1mg/ml | Invitrogen | 17104-019 |
| Trypsin | 0.25% | Invitrogen | 15090046 |
| Knockout Serum Replacement | 20% | Invitrogen | 10828010 |
| CaCl2 | 1mM | Sigma | C7902 |

**Table 3: Composition of neural induction media with growth factors.**

| Component | Volume (per 100ml of media) | Source | Catalogue number |
|---|---|---|---|
| DMEM-F12 | 50ml | Invitrogen | 10565-018 |
| Neurobasal media | 50ml | Invitrogen | 21103-049 |
| N2-suppliment (100X) | 0.5ml | Invitrogen | 17502048 |
| B27-suppliment (50x) | 1ml | Invitrogen | 17504044 |
| Glutamax (100X) | 0.5ml | Invitrogen | 35050-061 |
| Penn-Strep (100X) | 1ml | Invitrogen | 15070063 |
| EGF (20μg/ml) | 100μl | Sigma | 9644 |
| FGF (20μg/ml) | 100μl | Peprotech | 100-18B |
| Insulin (2mg/ml) | 250μl | Sigma | 16634 |

**Table 4: List of antibodies**

| Antibody | Source | Catalogue number | Dilution used |
|---|---|---|---|
| Twist1 | Abcam | Ab50887 | 1:100 |
| TFAP2a | Santacruz | Sc12726 | 1:100 |
| P75 | Promega | G323A | 1:500 |
| Sox1 | R&D | AF3369 | 1:100 |
| Nestin | Millipore | MAB5326 | 1:200 |
| Fibronectin (Fn1) | Millipore | AB2033 | 1:200 |
| Sox2 | Santacruz | Sc17320 | 1:100 |
| β-catenin | Abcam | Ab16051 | 1:250 |
| N-cadherin | BD biosciences | 610921 | 1:200 |

**Table 4: List of qPCR primers**

| Gene | Forward primer | Reverse primer |
|---|---|---|
| Id2 | GACAGCAAAGCACTGTGTGG | TCAGCACTTAAAAGATTCCGTG |
| Sox1 | TCAAGGAAACACAATCGCTG | ATTATTTTGCCCGTTTTCCC |
| Tfap2a | ATGCTTTGGAAATTGACGGA | ATTGACCTACAGTGCCCAGC |
| Sox10 | AGCTCAGCAAGACGCTGG | CTTTCTTGTGCTGCATACGG |
| Twist1 | TCCATTTTCTCCTTCTCTGGAA | GGCTCAGCTACGCCTTCTC |
